# CRISPRi with barcoded expression reporters dissects regulatory networks in human cells

**DOI:** 10.1101/2024.09.06.611573

**Authors:** Jinyoung Kim, Ryan Y. Muller, Eliana R. Bondra, Nicholas T. Ingolia

**Affiliations:** Department of Molecular and Cell Biology, University of California, Berkeley, Berkeley, CA 94720, USA; Center for Computational Biology, University of California, Berkeley, Berkeley, CA 94720, USA; California Institute for Quantitative Biosciences, University of California, Berkeley, Berkeley, CA 94720, USA

## Abstract

Genome-wide CRISPR screens have emerged as powerful tools for uncovering the genetic underpinnings of diverse biological processes. Incisive screens often depend on directly measuring molecular phenotypes, such as regulated gene expression changes, provoked by CRISPR-mediated genetic perturbations. Here, we provide quantitative measurements of transcriptional responses in human cells across genome-scale perturbation libraries by coupling CRISPR interference (CRISPRi) with barcoded expression reporter sequencing (CiBER-seq). To enable CiBER-seq in mammalian cells, we optimize the integration of highly complex, barcoded sgRNA libraries into a defined genomic context. CiBER-seq profiling of a nuclear factor kappa B (NF-κB) reporter delineates the canonical signaling cascade linking the transmembrane TNF-alpha receptor to inflammatory gene activation and highlights cell-type-specific factors in this response. Importantly, CiBER-seq relies solely on bulk RNA sequencing to capture the regulatory circuit driving this rapid transcriptional response. Our work demonstrates the accuracy of CiBER-seq and its potential for dissecting genetic networks in mammalian cells with superior time resolution.

## Introduction

CRISPR-based tools enable powerful reverse genetics approaches in mammalian cells that promise to reveal diverse biological pathways and regulatory mechanisms. CRISPR has been widely adopted due to its efficiency in perturbing genes and its flexibility in designing single guide RNAs (sgRNAs) that target specific sequences^1–3^. A critical challenge in any CRISPR screen is linking these sgRNAs with accurate and sensitive measurement of cellular phenotypes in high throughput. Survival and growth of genetically perturbed cells can be measured easily, but often provides an indirect and confounded view into specific pathways and processes. Fluorescent protein reporters can provide pathway-specific measurements^4^, but high-throughput fluorescence-based screening requires cell sorting, which imposes a bottleneck on cell population and discretizes reporter levels. Additionally, the limited time resolution of cell sorting makes it inadequate to study the fast kinetics of biological pathways. Single-cell sequencing of CRISPR-perturbed cell populations provides multidimensional and complex phenotypic readouts but can be costly and impractical to implement for studying one specific pathway or response^5^. Expressed RNA barcodes circumvent these limitations and have provided insights into transcriptional regulation by cis-acting sequences and transcription factors^6–9^. Many studies could benefit from direct RNA measurements of gene regulation phenotypes, made from bulk cell populations, that focus specifically on a pathway of interest.

To this end, our lab combined CRISPR interference (CRISPRi) with barcoded expression reporter sequencing (CiBER-seq) to measure cellular responses caused by sgRNA-mediated knockdown in budding yeast^10^. CiBER-seq couples a transcriptional reporter with a library of sgRNAs that perturb individual genes across the entire genome. Each sgRNA is linked to a specific nucleotide barcode that is embedded in the reporter^10^. The expressed RNA abundance of each barcode, measured by deep sequencing, reflects reporter activity levels in the cells expressing the associated sgRNA. A single CiBER-seq experiment captures the molecular phenotypes caused by genetic perturbations genome-wide in a single, bulk RNA sequencing library. Importantly, CiBER-seq and the similar ReporterSeq method overcome the limitations of single-cell sequencing and cell sorting, enabling the dissection of complex and fast-acting regulatory circuits^11^.

While CiBER-seq has been previously established in budding yeast, further adaptation is needed to carry out CiBER-seq screening in other organisms. Here, we extend CiBER-seq to mammalian cells. We adapt site-specific recombinases for highly efficient, single-copy integration of genome-wide barcoded sgRNA libraries into a human cell line. As proof of principle, we express RNA barcodes from two matched promoters and conduct CiBER-seq screening. In this situation, when no genes are expected to affect the promoters differentially, we rigorously control false discovery at the guide level and discover and correct for barcode-dependent effects, enabling more accurate measurement of gene knockdown phenotypes. We then use mammalian CiBER-seq to profile the genetic requirements for cytokine signaling to nuclear factor kappa B (NF-κB) and capture strong effects from disrupting the signaling pathway, from the transmembrane receptor through the transcription factor itself. Our system can be adapted to investigate the genetic regulators of almost any pathway in mammalian systems at a genome-wide scale.

## Results

### Single-copy genomic integration of dual barcoded CRISPRi sgRNA libraries

CiBER-seq measures how CRISPRi depletion or other CRISPR-mediated genetic perturbations affect a transcriptional reporter through deep sequencing of an expressed mRNA barcode **(Fig 1A)**^10^. Each sgRNA in a genome-wide library is linked to sgRNA-specific nucleotide barcodes on the same plasmid, embedded in a transcriptional reporter. The abundances of each barcode can be read out in parallel by deep sequencing, which provides a comprehensive profile of reporter expression across each individual sgRNA. The linkage between each RNA barcode and its associated sgRNA is critical to interpreting these data.

**Figure 1:**
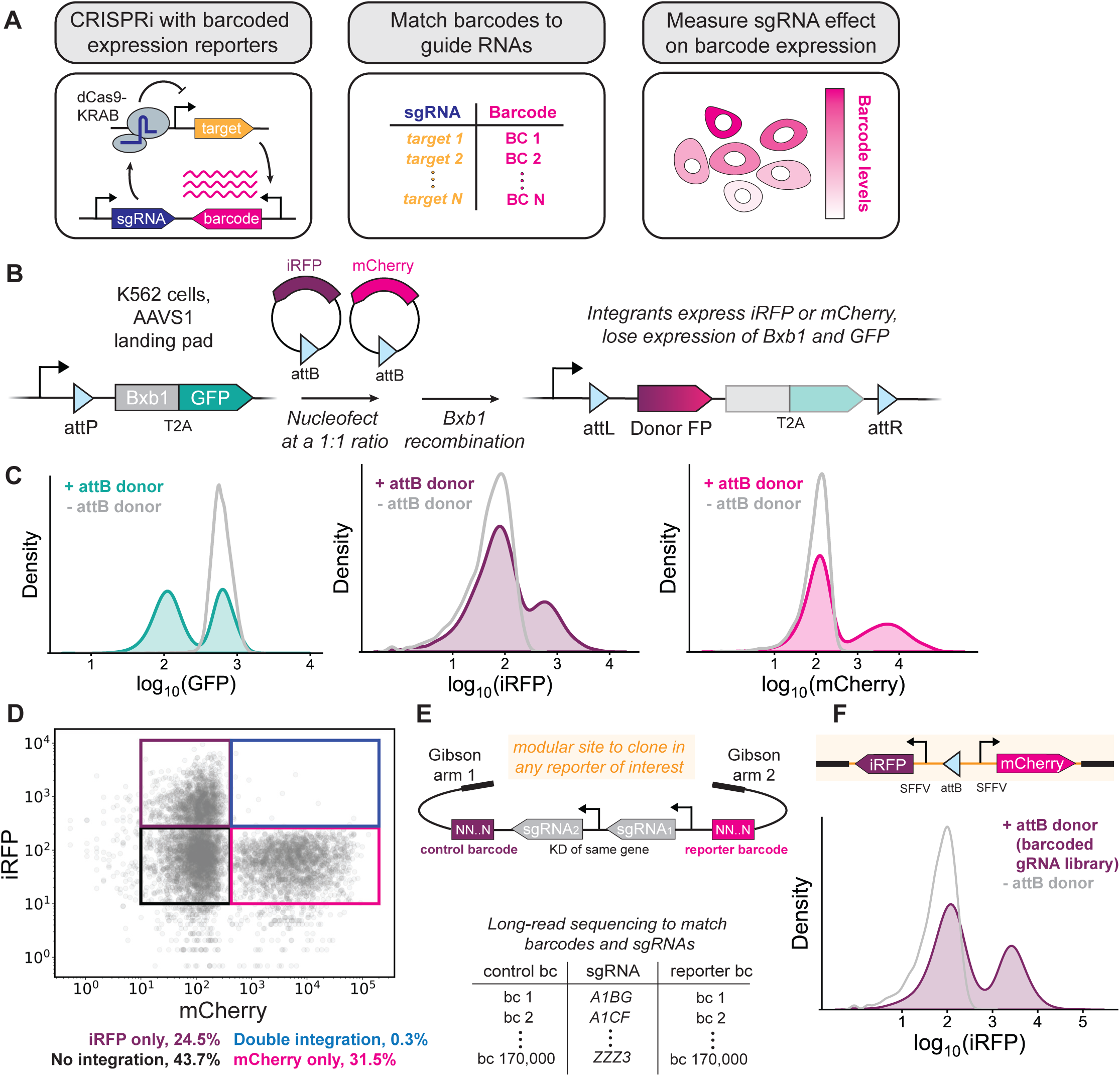
Single-copy genomic integration of dual barcoded sgRNA libraries. **(A)** CRISPRi with barcoded expression reporters. **(B)** Three-color flow cytometry assay to estimate Bxb1 integration efficiency and copy number. Cells that integrate an attB donor plasmid should lose expression of GFP and gain expression of either iRFP or mCherry. **(C, D)** Flow cytometry of cells (cJK1001) nucleofected with attB donor plasmids encoding either iRFP or mCherry. **(E)** Schematic of modular barcoded sgRNA plasmid library and corresponding lookup table generated by long-read sequencing. **(F)** Cells that integrate the barcoded sgRNA library express iRFP.

CRISPRi-based screens in human cells are commonly performed by lentiviral delivery of sgRNA libraries. However, lentiviral delivery has several limitations for barcode-based screens. Lentiviruses are pseudodiploid, and the reverse transcriptase that acts prior to integration is prone to template switching between these two genomes, resulting in scrambling between barcodes and sgRNAs^12–14^. Additionally, it is difficult to control the number of integration events per cell, and multiple sgRNAs may be introduced into the same cell, adding experimental noise^15^. Lentiviruses also integrate randomly in the genome, which may lead to variable expression of transcriptional reporters.

To overcome these limitations, we developed a strategy for site-specific integration of barcoded expression reporters and sgRNAs at a single, defined locus in human cells using the serine recombinase Bxb1. Bxb1 recombinase has been previously utilized for site-specific integration of complex libraries—including sgRNA libraries—in human cells^16,17^. We introduced a Bxb1 “landing pad” into the *AAVS1* safe-harbor locus^18^. The landing pad contained a constitutive promoter upstream of an attP recombinase site, followed by a gene encoding the Bxb1 recombinase fused with EGFP **(Fig 1B)**. This landing pad expresses Bxb1 constitutively prior to recombination but stops expressing Bxb1 after recombination **(Fig 1B)**. Starting with K562 cell lines that express the CRISPRi effector protein dCas9-KRAB either constitutively or from a doxycycline-inducible promoter, we created clonal derivatives with our attP landing pad **(Fig S1)**. We also over-expressed Bxb1 recombinase elsewhere in the genome to increase recombination efficiency.

To measure the efficiency of Bxb1-mediated integration and the number of integration events per cell, we developed a three-color flow cytometry assay **(Fig 1B)**. We created donor plasmids containing attB recombinase sites next to a fluorescent protein gene lacking a promoter. Integration of these donor plasmids into the landing pad should enable expression of the fluorescent protein, driven by the promoter in the landing pad, while simultaneously shutting down EGFP expression. We delivered an equal mixture of iRFP and mCherry donor plasmids by nucleofection and saw that over half of cells lost EGFP expression and gained either iRFP or mCherry expression **(Fig 1C)**. Roughly equal numbers of cells were positive for exactly one of the iRFP or mCherry markers, while only a tiny fraction of cells appeared positive for both, indicating that most cells only had a single integration event **(Fig 1D)**. We concluded that Bxb1 recombination overcomes the limitations of lentiviral delivery while maintaining high efficiency, single-copy integration.

We next constructed a donor library of barcoded sgRNA libraries suitable for CiBER-seq screening. We prepared a 21,554-member plasmid pool based on a recent compact, genome-wide dual-sgRNA CRISPRi library that targets each human gene with a matched pair of sgRNAs against that gene, delivered together **(Fig 1E)**^19^. These sgRNAs were chosen as the best two sgRNAs for the gene, based on aggregated empirical activity data and computational predictions^19^. We then introduced two unique random 25 nucleotide barcodes into these plasmids and ascertained the linkage between barcodes and sgRNAs on each plasmid by long-read sequencing, thus generating a lookup table that could later be used to link barcode abundance to sgRNA identity. Our library contained an average of 8 barcodes per sgRNA, providing substantial internal replication for each sgRNA.

We tested integration of our donor library using fluorescent protein reporters. This modular library contained a short spacer between the two barcodes where transcriptional reporters of interest can be inserted for CiBER-seq screening^20^. We introduced mCherry and iRFP, each expressed from identical constitutive promoters, along with an attB donor site, into this spacer **(Fig 1F, S1B)**. Nucleofection of our human CiBER-seq plasmid library yielded mCherry and iRFP expression in 29% of cells in the absence of any selection or screening for integrants, indicating that Bxb1 integrated our plasmid library robustly and validating our strategy for large-scale, barcode-based genetic screens in human cells.

### Matched barcoded expression reporters in mammalian CiBER-seq

To validate our mammalian cell CiBER-seq strategy and establish a baseline for its sensitivity, we conducted a CiBER-seq screen using barcoded reporters driven from identical, constitutive promoters. These matched promoters should yield the same expression level across the paired barcodes, and thus statistically indistinguishable read counts upon deep sequencing. Noise and sequence-dependent bias could result in apparent differential expression between paired barcodes, however, given the high quantitative sensitivity of CiBER-seq^21^. We wished to measure the magnitude of these effects, which could confound our interpretation of differential effects on transcriptional reporters.

We thus created a “dual-SFFV” CiBER-seq library containing iRFP and mCherry, each expressed from the strong and constitutive SFFV promoter **(Fig 2A)**. We nucleofected this dual-SFFV library into our K562 cell line containing inducible dCas9-KRAB and the attB landing pad in biological triplicate and selected for cells that integrated the dual-SFFV plasmid library with puromycin **(Fig 2B)**. As expected, flow cytometry indicated that over 99% of cells robustly expressed both iRFP and mCherry after 1 week of selection **(Fig 2C)**. We induced CRISPRi knockdown for 72 hours by adding doxycycline, and then collected both uninduced and induced cells. Following RNA extraction, we generated a barcode sequencing library by reverse transcription with gene-specific primers that contained unique molecular identifiers (UMIs), allowing us to distinguish barcodes that originated from different mRNA molecules. We sequenced barcodes and UMIs, removed PCR duplicates, and performed DESeq2 analysis to test for differences across samples and between paired mCherry and iRFP barcodes^22^. None of the 167,916 unique barcodes showed a significant difference in abundance between CRISPRi-uninduced and induced samples, indicating that there is little to no noise introduced from library preparation for near-replicate samples **(Fig S2A, S2B)**. The strong concordance suggests that inducible CRISPRi systems enable high-precision CiBER-seq measurements that inherently control many technical effects, as we saw in yeast^10,23^.

**Figure 2:**
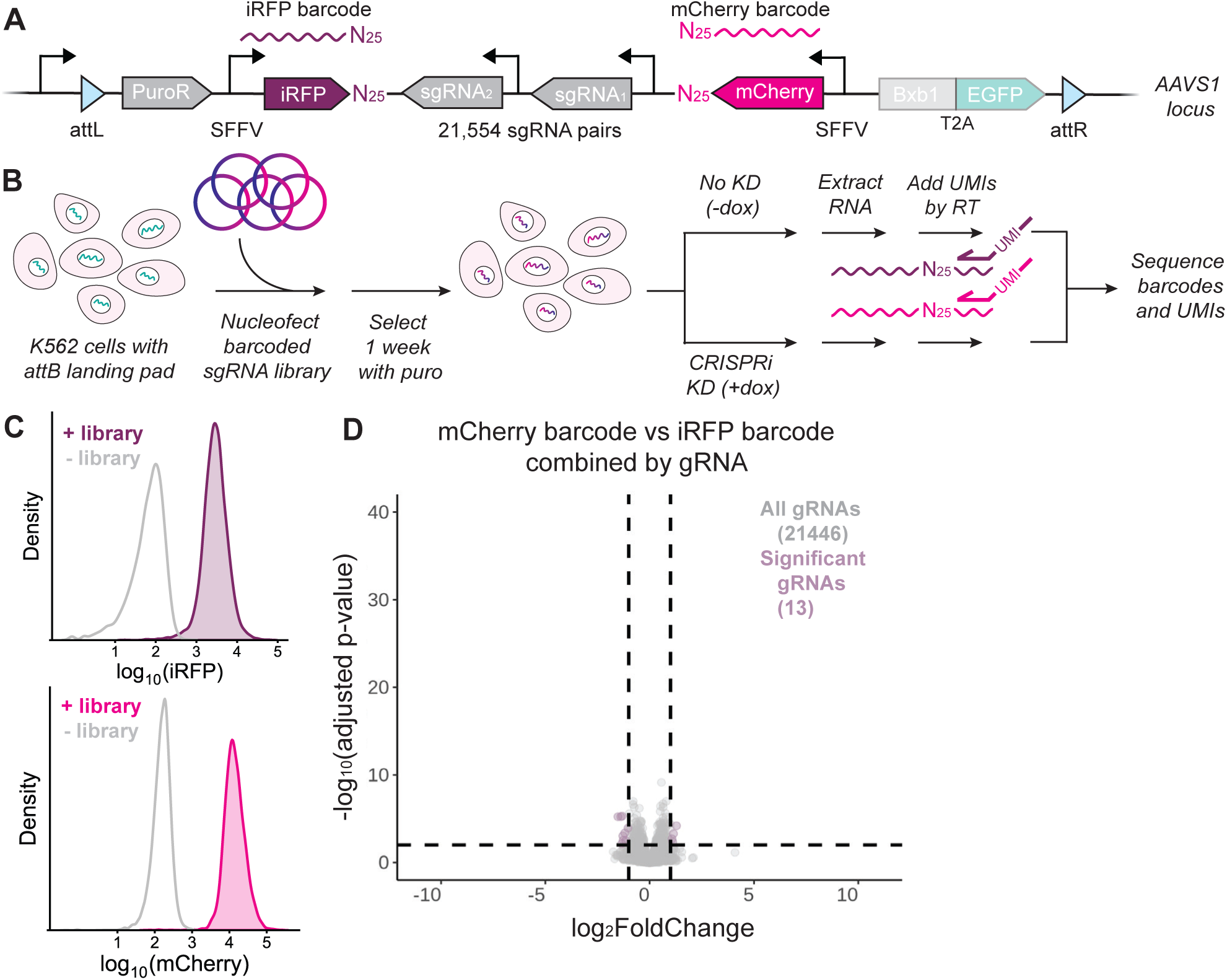
Matched barcoded expression reporters in mammalian CiBER-seq. **(A)** Schematic of the dual SFFV library integrated into cells at the landing pad locus. **(B)** General schematic for integrating the barcoded sgRNA library into cells and preparing libraries from harvested cells. **(C)** Cells selected for dual SFFV library integration express iRFP and mCherry uniformly. **(D)** DESeq2 analysis of mCherry vs iRFP barcodes (from both the pre- and post-CRISPRi induction samples) in the dual SFFV CiBER-seq screen. Each point represents a sgRNA with its barcodes combined. Dashed lines represent adjusted p-value < 0.01 and log2-fold change > 1 or < -1.

We did see some evidence for differences between the two barcodes in a matched pair **(Fig S2C-E)**. Although most barcodes did not seem to have a strong sequence-dependent effect, a minority of barcode pairs were differentially expressed up to 8-fold. Some of the barcode-dependent noise was mitigated by removal of PCR duplicates using UMI collapse **(Fig S2C-E)**. Most of the barcode-dependent noise was ameliorated when the barcodes were combined by sgRNA, resulting in only 13 out of 21446 sgRNAs that had a weak but marginally significant effect by DESeq2 **(Fig 2D)**. Although combining barcodes largely mitigated these barcode-specific effects, we reasoned that data from our dual-SFFV screen can be used to normalize barcodes in any future CiBER-seq screens. This built-in normalization could be especially useful for CiBER-seq screens that have lower coverage and fewer barcodes per sgRNA represented, where barcode-dependent effects may appear particularly strong.

### Mammalian CiBER-seq identifies regulators of NF-κB in response to TNF-α stimulation

We next sought to demonstrate how mammalian CiBER-seq could dissect signaling pathways using a downstream transcriptional reporter. Among other advantages, the RNA reporters used in CiBER-seq promise higher time resolution than fluorescent protein reporters, which must be exported to the cytoplasm, undergo translation, fold, mature, and accumulate to detectable levels before measurement. Cell signaling pathways operate quickly, and many transcription factors are induced within minutes of receiving a stimulus^24^. For example, the pro-inflammatory NF-κB transcription factor is activated within half an hour of cytokine stimulation, and we confirmed that the known target gene *NFKBIA* was strongly induced within 1 hour of tumor necrosis factor-alpha (TNF-α) stimulation **(Fig S3A)**^25–27^. To assay NF-κB activity, we designed a synthetic reporter with 3⨉ NF-κB binding motifs upstream of a minimal promoter driving mCherry. We then integrated this NF-κB reporter, along with constitutive iRFP, into our K562 landing pad cell line. Levels of reporter RNA were elevated within 1 hour of TNF-α stimulation, matching the speed of endogenous gene activation, and continued to accumulate over the next several hours **(Fig 3A)**. In contrast, matched flow cytometry indicated that mCherry fluorescence only increased detectably after 3 hours and was not clearly separated from background until 5 hours after TNF-α treatment **(Fig 3B)**. Thus, we chose to conduct our mammalian CiBER-seq screen at an earlier 2-hour timepoint, when fluorescence was not visible, and a later, 5-hour timepoint.

**Figure 3:**
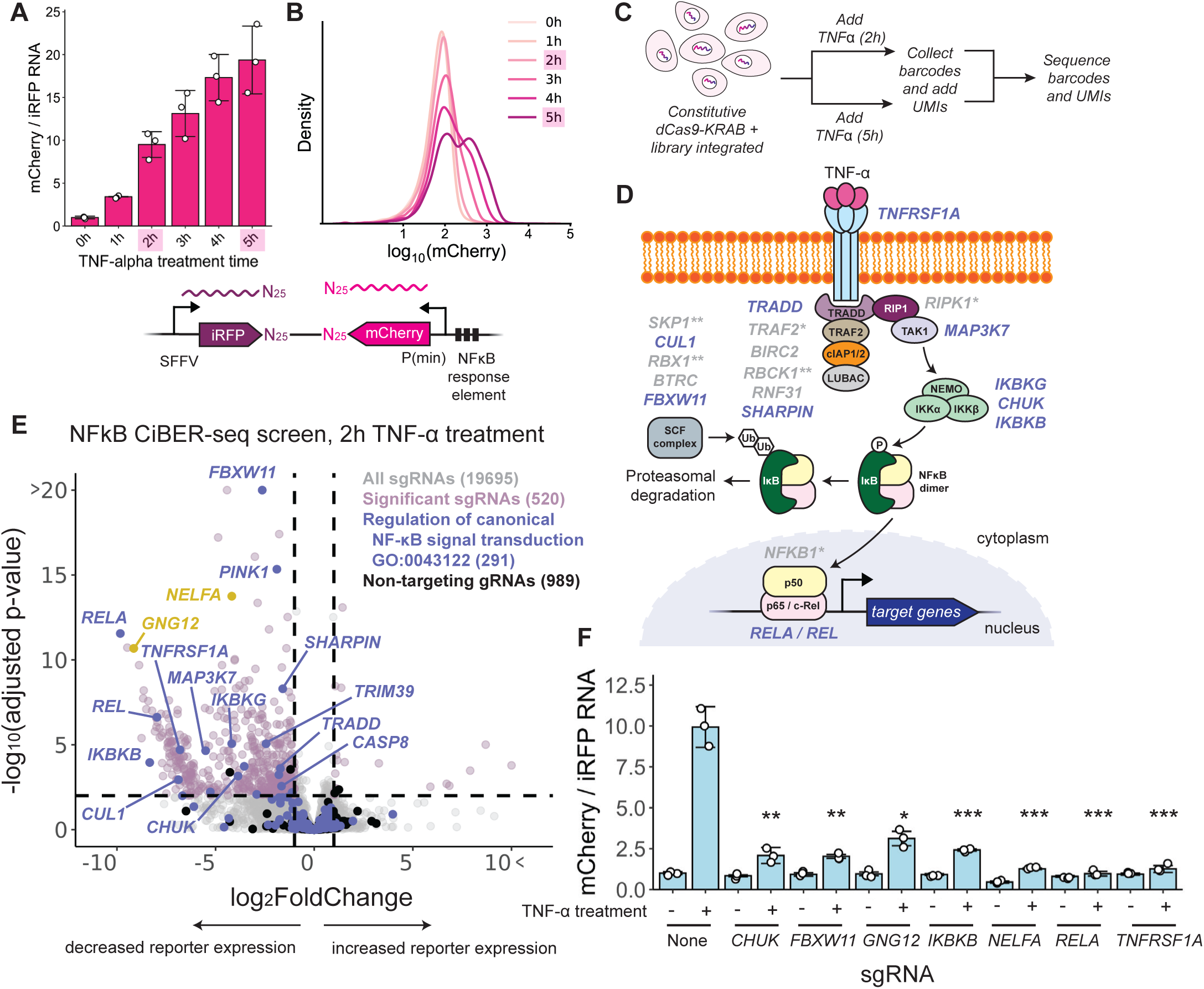
Mammalian CiBER-seq identifies regulators of NF-κB in response to TNF-α stimulation. **(A)** RT-qPCR of the mCherry NF-κB reporter normalized to constitutive iRFP across different timepoints of TNF-α stimulation (n=3). **(B)** As in (A), but mCherry analyzed by flow cytometry. **(C)** Schematic of NF-κB CiBER-seq experiment, where TNF-α was added at two different timepoints to cells with the NF-κB barcoded sgRNA library integrated. **(D)** Proteins involved in NF-κB pathway activated by TNF-α, with human gene names displayed. Genes highlighted in purple were significant in E. * indicates genes where knockdown is not expected to prevent NF-κB activation based on previous literature. ** indicates genes where the sgRNA dropped out in the screen. **(E)** DESeq2 analysis of genome-wide CiBER-seq screen for regulators of NF-κB. Each point represents a single sgRNA. Dashed lines represent adjusted p-value < 0.01 and log2-fold change > 1 or < -1. **(F)** RT-qPCR of the mCherry NF-κB reporter normalized to constitutive iRFP at 2 hrs of TNF-alpha stimulation in cells expressing different sgRNAs (n=3). Statistical significance was calculated by two-sided t tests between the no-sgRNA sample and each individual sgRNA samples at 2 hrs of TNF-α treatment. * indicates p-value < 0.001, ** indicates p-value < 0.00001, and *** indicates p-value < 0.000001.

We then conducted a screen for regulators of NF-κB, using a CiBER-seq library with the NF-κB reporter. We first introduced this library into the landing pad in our K562 cell line expressing constitutive dCas9-KRAB and selected integrants. We then treated cells with TNF-α for either 2 or 5 hours, harvested RNA, and sequenced barcodes in each sample **(Fig 3C)**. We first confirmed that constitutive iRFP barcode counts between timepoints within a replicate correlated well (Spearman’s ρ = 0.992-0.993), indicating that library preparation was robust **(Fig S3B)**. In contrast, we saw many NF-κB reporter barcodes with low abundance relative to the matched constitutive partner, indicating that cells containing that barcode showed weaker NF-κB induction in response to TNF-alpha **(Fig S3C)**. Indeed, independent DESeq2 analysis on each TNF-alpha induction timepoint, normalized to dual SFFV data, revealed hundreds of sgRNAs that blocked NF-κB reporter activation, whereas the vast majority of our 989 non-targeting sgRNAs had no significant effect. We thus wanted to map these sgRNAs onto the signaling pathway that transduces extracellular TNF-α binding to NF-κB transcriptional responses **(Fig 3D)**.

Strikingly, sgRNAs targeting *RELA*—which encodes a subunit of the NF-κB transcription factor that contains a transactivation domain—showed the strongest effect in our screen **(Fig 3E)**^28^. The *REL* gene, encoding another transcription-activating subunit of NF-κB, was also amongst the top hits^28^. We identified many other components of the TNF-α signaling cascade, starting with the TNF-α receptor 1 (*TNFRSF1A*) itself and extending through factors that are recruited to the receptor (*TRADD*, *SHARPIN*, *MAP3K7*) upon cytokine binding^25,29^. Not every component of the signaling pathway reduced the mCherry to iRFP ratio, likely because knockdown of a single gene serving as an intermediary adapter cannot prevent signaling in some cases. In unstimulated conditions, NF-κB activity is restrained by the inhibitor of kappaB (IκB); after TNF-α treatment, IκB is phosphorylated by IKK, ubiquitinated by SCF-βTrCP, and degraded by the 26S proteasome^25^. Knockdown of every component of the IKK complex (*IKBKG*, *CHUK*, *IKBKB*) led to a significant reduction in the mCherry to iRFP ratio. Additionally, sgRNAs targeting some components of the SCF complex (*CUL1*, *FBXW11*) involved in degrading IκB also reduced the mCherry to iRFP ratio – notably, the βTrCP2 (*FBXW11*) sgRNA was the second most significant hit in our screen^30^. More broadly, gene ontology (GO) analysis of targets that decreased the mCherry to iRFP barcode ratio upon 2 hours of TNF-alpha induction revealed annotations for NF-κB signal transduction **(Fig S3F)**^31^. Analysis of the 5-hour TNF-α induction timepoint revealed a similar spectrum of phenotypes **(Fig S3D)**. Indeed, direct comparison between the 2-hour and the 5-hour induction timepoints identified no significant sgRNAs, indicating that the CiBER-seq screen yielded clear results at this early timepoint **(Fig S3E)**. Interestingly, there were very few sgRNAs that further increased the mCherry to iRFP barcode ratio at either timepoint, and GO analysis of these sgRNAs yielded no results.

We selected several sgRNAs that had a significant reduction in their mCherry to iRFP barcode ratios for individual validation. Integration and expression of the selected sgRNAs led to strong knockdown of their respective target genes with efficiencies ranging from 70-98% **(Fig S3G)**. Knockdown of some of the canonical components of NF-κB signaling blocked induction of the mCherry reporter when cells were treated with TNF-α, consistent with screen results **(Fig 3F)**. Surprisingly, knockdown of two non-canonical signaling components, negative elongation factor subunit A (*NELFA*) and guanine nucleotide-binding protein subunit gamma-12 (*GNG12*), also both blocked induction of the mCherry reporter in the same experiment **(Fig 3F)**. Negative elongation factor stabilizes paused RNA polymerase II to allow for rapid gene expression changes, so the role of this factor was consistent with our understanding of NF-κB-mediated gene regulation^32^. Although it is not part of the canonical pathway, *GNG12* has been linked to NF-κB pathway activation in cancer^33,34^, so it may play a role in NF-κB activation in K562 cells. These results demonstrate how mammalian CiBER-seq can reveal key components of a biological pathway with fast kinetics.

## Discussion

In this study, we develop mammalian CiBER-seq, a genome-wide CRISPRi screening platform with an mRNA barcode phenotypic readout that allows us to dissect signaling pathways in human cells. We overcome challenges to barcoded CRISPR screening by recombinase mediated integration. As proof of principle, we show that mRNA barcodes can be robustly read out by deep sequencing and use this data to correct for barcode-dependent effects in any future screens. Finally, we conduct a CiBER-seq screen that recapitulates the known pathway of NF-κB activation in response to cytokine signaling and identifies non-canonical factors that may have cell-type-specific roles.

RNA barcodes in CiBER-seq promise advantages for diverse applications compared to other screenable reporters. In this work, we demonstrated that CiBER-seq can address rapid regulatory responses, because mRNA expression precedes protein expression, and it accumulates to higher levels more quickly. Cells sense diverse changes in their environment within seconds and rapidly alter the expression of many genes in response; CiBER-seq will be useful for studying these fast-acting pathways^35^. Furthermore, CiBER-seq and similar approaches that utilize RNA expression yield clear measurements of phenotypes and genetic perturbations where protein synthesis may be limited, as shown in yeast^10,11,36^. In contrast, translation-limiting conditions can impair synthesis of fluorescent reporter proteins. These advantages extend to reporters where functional protein output is naturally low, such as nonsense-mediated decay reporters containing premature termination codons, where the reporter protein is difficult to observe by flow cytometry^23,37,38^. More broadly, mammalian CiBER-seq has the potential to be a tool for dissecting the regulation of lowly expressed endogenous genes where changes in fluorescence may be difficult to distinguish.

In this work, we established the fundamental framework for performing CiBER-seq in mammalian cells. This core can be extended to study more complex phenotypes. While we measured a single reporter barcode along with a control barcode for each sgRNA, adding more barcodes would create multiplexed reporters that capture several molecular phenotypes in a single experiment. Conversely, because CiBER-seq sample collection and library preparation are very straightforward, it should be possible to challenge cells with many environmental conditions and drugs in parallel in a single experiment^11^. High coverage in these experiments might be aided by barcoded sgRNA sub-libraries designed for more targeted CiBER-seq. Sampling multiplexed reporters, treatments, and dense time-courses should provide high-dimensional quantitative phenotypes for each guide RNA. We have also shown how CiBER-seq can measure translational or post-translational regulation using reporters driven by synthetic transcription factors further expanding the range of biological processes it can address^10,23^. CiBER-seq has already led to many new biological insights in yeast. Its wide applicability and unique advantages position CiBER-seq well to dissect the many complex genetic networks in mammalian cells.

## Author contributions

J.K., R.Y.M., and N.T.I. designed the overall strategy. J.K. designed and conducted experiments based on preliminary work from R.Y.M. with help from E.R.B. J.K. analyzed the data with guidance from N.T.I. J.K. and N.T.I. wrote the manuscript with input from R.Y.M. and E.R.B. All authors discussed the results, interpretation, and contributed to the final manuscript.

## Declaration of interests

N.T.I. holds equity in Velia Therapeutics and holds equity and serves as a scientific advisor to Tevard Biosciences.

## Acknowledgements

We thank Joseph Lobel, J. Wren Kim, and all members of the Ingolia lab for thoughtful discussion and commentary. We also thank the Flow Cytometry Core Facility, the Tissue Culture Facility, and the DNA Sequencing Facility at the University of California, Berkeley for their guidance. This work was supported by the National Institute of General Medical Sciences of the National Institutes of Health under award numbers R01 GM135233 (N.T.I.) and R35 GM139488 (E.R.B), by the Shurl & Kay Curci Foundation (N.T.I.), and by the National Science Foundation Graduate Research Fellowship under Grant No. DGE 1752814 (J.K.). Any opinions, findings, and conclusions or recommendations expressed in this material are those of the authors and do not necessarily reflect the views of the National Science Foundation. Short-read sequencing was performed at the UCSF CAT, supported by UCSF PBBR, RRP IMIA, and NIH 1S10OD028511-01 grants. Long read sequencing was carried out at the DNA Technologies and Expression Analysis Cores at the UC Davis Genome Center, supported by NIH Shared Instrumentation Grant 1S10OD010786-01.

## Materials and Methods

### Plasmid construction

All plasmids used are listed in Table S1, and all oligos are listed in Table S2. All plasmids were generated by standard restriction digestion, PCR amplification using Q5 polymerase, and Gibson assembly. Whole plasmid sequencing using Oxford Nanopore Technology with custom analysis and annotation (Plasmidsaurus) was used to verify all constructs.

### Genome-wide barcoded sgRNA plasmid library construction and sequencing

The genome-wide barcoded sgRNA library was cloned in a 2-step process. The first step involved inserting a library of 21,554 human sgRNAs into a base vector. pJK054A, the base vector, was amplified using oJK405 and oJK430. A plasmid library containing the top two sgRNAs for every human gene (dJR072) was a gift from Jonathan Weissman (Addgene, 187246). dJR072 was digested with three separate pairs of restriction enzymes to excise the tandem sgRNA cassette, to avoid the loss of sgRNAs cut by any individual enzyme pair and ensure that all sgRNAs would be represented in the final library. dJR072 was digested with 1) XbaI and BamHI-HF, 2) BsrBI and NotI-HF, or 3) ApaI and NotI-HF. Three different 100 μL Gibson reactions (NEBuilder HiFi, NEB, E2621) were prepared with 125 ng of the pJK054A PCR product and 1.25 ug of each dJR072 restriction digest and incubated at 50°C for 1 hr. The Gibson reactions were purified by spin column and eluted in 6 μL water. 1 μL of each Gibson reaction was electroporated into 20 μL MegaX DH10B T1^R^ Electrocomp cells (ThermoFisher, C640003) according to the manufacturer’s protocol. Cells were shaken at 37°C for 1 hr in recovery media then transferred into 100 mL LB+Chloramphenicol (Cam) and grown overnight at 28°C to OD_600_ = 2.0. Library coverage was estimated by serial dilution of transformants post-recovery and prior to overnight growth. At least 1000 unique transformants for each sgRNA in the library were obtained. The plasmid libraries were then purified from overnight cultures by maxiprep (Qiagen, 12662) and confirmed by whole plasmid sequencing and Sanger sequencing. Equal amounts of DNA from the three libraries were mixed, resulting in pLJK05.

The second step of cloning involved inserting random 25-nucleotide barcodes into pLJK05 in a 4-piece Gibson assembly reaction. 6 μg of pLJK05 was digested by SbfI-HF and purified using a spin column. The ampicillin resistance marker (AmpR) was amplified using oJK442 and oJK443 and purified by agarose gel electrophoresis followed by extraction of the product band. To insert barcodes, 250 ng of pLJK05 vector and 90 ng of AmpR were assembled with oJK441 and oJK444 at a 1:1:100:100 ratio. oJK0441 and oJK444 both contain random 25-nucleotide sequences in between the 25-nucleotide Gibson overhangs on either side. The 50 μL Gibson assembly was incubated at 50°C for 1 hr, purified by spin column, and eluted in 6 μL water. 0.5 μL of the Gibson assembly was transformed into 20 μL MegaX DH10B T1^R^ Electrocomp cells and grown at 37°C for 1 hr in recovery media. Dilutions of the transformation were plated to estimate the total number of transformants. The transformation was inoculated at several dilutions into LB+Cam+Carb media and grown overnight at 28°C to OD_600_ = 2.0. Liquid culture corresponding to approximately 170,000 transformants was harvested. Plasmid libraries were purified by maxiprep and confirmed by whole plasmid sequencing and Sanger sequencing, resulting in pLJK06. We found that introducing barcodes by assembly of single-stranded oligonucleotides avoided scrambling between barcode pairs that we attribute to heteroduplex formation and use this approach despite the relatively low efficiency of assembly reactions with single-stranded templates.

To assign barcodes to sgRNAs, long-read sequencing was performed. pLJK06 was digested with I-CeuI and purified by spin column. The resulting linearized plasmid library was sequenced on the PacBio Revio instrument. The barcode to sgRNA assignment was conducted using custom scripts.

To insert reporters into the modular barcoded sgRNA library, pLJK06 was linearized with PacI and plasmids containing reporters (pJKEB01 or pJK013D) were linearized with BciVI. These plasmids contain compatible Gibson homology arms next to their restriction sites. The digestions were purified by spin column and the library was produced by Gibson assembly. The Gibson reaction was purified, electroporated into MegaX DH10 T1^R^ Electrocomp cells, and purified from culture by maxiprep as described above, always ensuring >100 unique transformants per barcode. The resulting libraries were named pLJK08 (dual SFFV reporters) and pLJK07 (NF-κB reporter with SFFV control reporter).

### Cell culture

Human K562 and HEK293T Lenti-X cells were obtained from the UC Berkeley Cell Culture Facility. A polyclonal K562 CRISPRi dCas9-HA-BFP-KRAB cell line was gifted by Jonathan S. Weissman^3^. K562 cells were grown in RPMI 1640 with L-glutamine (Thermo Fisher Scientific) supplemented with 10% FBS, 1% sodium pyruvate, 100 units/mL penicillin, and 100 mg/mL streptomycin. K562 cells were maintained at a density of 0.5 x 10^6^ cells/mL. Human HEK293T Lenti-X cells were grown in DMEM + GlutaMAX (Thermo Fisher Scientific) supplemented with 10% FBS. All cell lines were grown at 37°C and 5% CO_2_. Cell lines were confirmed to be free of mycoplasma contamination (Bulldog Bio, 2523448).

### Generation of CiBER-seq cell lines

All cell lines used are listed in Table S3. To generate the K562 constitutive CRISPRi CiBER-seq cell line (cJK1003), the attP landing pad along with a cassette expressing Bxb1 and EGFP was integrated by Cas9 into the *AAVS1* locus of K562 CRISPRi dCas9-KRAB cells by nucleofecting pJK025B with pX330 (Addgene, 42230). Cells were selected using 250 μg/mL hygromycin (InvivoGen). A Bxb1 expression construct (pJK040A) was stably integrated at multiple copies into this cell line by Sleeping Beauty transposition then selected using 800 μg/mL G418 (InvivoGen). Monoclonal cell lines were isolated by sorting for cells expressing BFP and GFP and single-cell deposition.

To generate the K562 inducible CRISPRi CiBER-seq cell lines (cJK1001 and cJK1002), pTet-On Advanced (Taraka Bio, 630930) expressing the reverseTet-controlled transactivator protein (rtTA) and a plasmid encoding dCas9-KRAB driven from the Tet-On 3G promoter, both gifted to us by James Nuñez, were integrated into cells by lentiviral transduction with MOI > 1. The attP-Bxb1-EGFP cassette was subsequently introduced into cells by Cas9 as described above. Bxb1 expression constructs (pJK040A and pJK062A) were integrated into cells by Sleeping Beauty transposition then selected using 800 μg/mL G418 and 10 μg/mL blasticidin (InvivoGen). Monoclonal cell lines were isolated by sorting for cells constitutively expressing GFP and expressing BFP upon treatment with 1 μg/mL doxycycline (Sigma-Aldrich, D9891) for 48 hrs. cJK1001 has two copies and cJK1002 has one copy of the attP landing pad.

### Flow cytometry

To harvest cells for flow cytometry, cells were collected by centrifugation at 300 g for 5 min at RT, washed once in PBS, and resuspended in PBS supplemented by 1% FBS, and collected in tubes with cell strainer caps. All analyses were performed on an LSR Fortessa Analyzer (BD Biosciences). Cells were gated on forward scatter (FSC) and side scatter (SSC), and positive events were determined by non-fluorescent negative control cells. Fluorescence was detected on the Pacific Blue (450/50 nm), FITC (530/30 nm), PE-Texas Red (610/20 nm), and APC-Cy7 (780/60 nm) channels. Flow cytometry data was analyzed using *FlowCytometryTools* in python.

For the three-color Bxb1 integration assay, 200,000 cells were nucleofected with either 500 ng pJK028A and 500 ng pJK029A or 1 μg pJKEB01 using the Lonza SF Cell Line 4D-Nucleofector X Kit S (Lonza, V4XC-2032) according to manufacturer’s instructions. Cells were passaged for 10 days post-nucleofection to dilute out unintegrated plasmid and subjected to flow cytometry as described above.

### RT-qPCR

Cells were harvested for RT-qPCR by centrifugation and lysed in TRI reagent (Sigma-Aldrich, T9424). To isolate RNA, 200 μL chloroform per 1 mL TRI reagent was added, shaken vigorously, incubated at RT for 5 min, then centrifuged at 12,000 g for 15 min at 4°C. The aqueous phase was transferred to a new tube, and 1 μL GlycoBlue (ThermoFisher, AM9515) and 500 μL 2-propanol per 1 mL TRI reagent was added. Samples were incubated at RT for 10 min then centrifuged at 12,000 g for 10 min at 4°C. The supernatant was removed from the RNA pellet, and the pellet was washed by adding at least 1 mL 75% ethanol. Samples were vortexed and centrifuged at 7500 g for 5 min at 4°C. The RNA pellet was briefly dried and then resuspended in water. To synthesize cDNA, 1 μg total RNA was reverse transcribed using dT priming with Protoscript II (NEB, M0368L). qPCR was conducted using DyNAmo qPCR mastermix (ThermoFisher, F410L) according to manufacturer protocols using the Agilent Stratagene Mx3000P qPCR System. In addition to 3 biological replicates, 2-3 technical replicates were performed for all qPCR reactions. All primers (listed in Table S4) used for gene expression analysis were validated using a cDNA dilution standard curve.

### Western blotting

To perform western blotting, cells were harvested by centrifugation, washed with PBS, and lysed with buffer (140 mM KCl, 10 mM HEPES, 5 mM MgCl_2_, 1 % Triton X-100, 1 mM TCEP, 2 U/μL Turbo DNase) on ice for 20 min. Lysates were clarified by centrifugation at 20,000 g for 10 min at 4°C. The Pierce 660nm assay was performed to determine protein concentration. Equal amounts of total protein were run on a 4 to 12% polyacrylamide Bis-Tris gel (ThermoFisher Scientific, NW04125BOX) then transferred to a nitrocellulose membrane. Membranes were blocked with 5% milk in TBST (0.1% Tween-20) for 1 hr at RT. Primary antibodies were incubated overnight at 4°C, and secondary antibodies were incubated for 1 hr at RT. Membranes were washed three times with TBST between each step. The antibodies and their corresponding dilutions used in this study are: β-Actin (D6A8) Rabbit mAb HRP Conjugate (Cell Signaling Technology, 12620) at a 1:2000 dilution; V5-Tag (D3H8Q) Rabbit mAb (Cell Signaling Technology, 13202) at a 1:1000 dilution; HA-Tag (C29F4) Rabbit mAb (Cell Signaling Technology, 3724) at a 1:1000 dilution; Anti-rabbit IgG, HRP-linked Antibody (Cell Signaling Technology, 7074) at a 1:3000 dilution. Blots were developed with Pierce ECL Western Blotting Substrate (ThermoFisher Scientific, 32106) and visualized by a FluorChem R imaging system.

### NF-κB TNF-alpha induction time-course

To conduct the NF-κB time-course, a stable cell line cJK068 was first created by nucleofecting 1 μg pJK013D into cJK1003 using the Lonza SF Cell Line 4D-Nucleofector X Kit S according to manufacturer’s instructions. Cells with pJK013D stably integrated at the Bxb1 landing pad were selected for 14 days with 1 μg/mL puromycin (InvivoGen). cJK068 was treated with 5 ng/mL TNF-alpha (Sigma-Aldrich, H8916) for 1 to 5 hrs in triplicate. Each sample was split in half and collected for flow cytometry and RT-qPCR as described above.

### CiBER-seq screening in cells

CiBER-seq screening was carried out in cJK1001, cJK1002, or cJK1003 in biological triplicate. Approximately 12 million cells per replicate were spun down and resuspended in nucleofection solution from the Lonza SF Cell Line 4D-Nucleofector X Kit L (Lonza, V4XC-2012) and nucleofected in 12 reactions with 5 μg pLJK07 or pLJK08 per 1 million cells according to manufacturer’s instructions. Cells were recovered in media for 72 hrs and then cells that integrated the plasmid library by Bxb1 recombination were selected with 1 μg/mL puromycin (InvivoGen) for 7 days. Cells in selection were expanded and maintained at >500x coverage per barcode and >4000x coverage per sgRNA. For the dual SFFV screen **(Fig 2)**, cells were then split in half and resuspended in puro-free media with or without 1 μg/mL doxycycline. Cells were collected in TRI reagent 72 hrs post induction. For the NF-κB screen **(Fig 3)**, cells were then split in thirds and resuspended in puro-free media without TNF-alpha, with 5 ng/mL TNF-alpha for 2 hrs, or with 5 ng/mL TNF-alpha for 5 hrs. Cells were promptly collected in TRI reagent.

### Barcode isolation, library preparation, and deep sequencing

To isolate barcodes, RNA was harvested from the TRI reagent lysate as described in the RT-qPCR section. 400 μg of RNA was treated with Turbo DNase (ThermoFisher Scientific, AM2238) for 30 min at 37°C. RNA was recovered by adding 3 volumes of TRI reagent and extracting as described above. For each sample, 100 μg of RNA was reverse transcribed using Protoscript II and two gene-specific primers at a 1:1 ratio to add UMIs (oJK509, oJK510). cDNA was treated with 1 μL each of RNase A and RNase H (NEB M0297S and ThermoFisher, EN0531) for 30 min at 37°C. cDNA from each sample was purified by spin column then quantified by qPCR as described in the RT-qPCR section. cDNA was then split in half and used as input for two step-1 PCR reactions. oJK292 and oJK547 were used to amplify the iRFP barcode whereas oJK481 and oJK548 were used to amplify the mCherry barcode. Both step-1 PCR reactions were performed in 50 μL with a 98°C initial denaturation for 30 s, followed by 6 cycles of 98°C for 10 s, 68°C for 15 s, and 72°C for 15 s, with a final extension of 72°C for 2 min using Q5 Hot Start High-Fidelity DNA Polymerase (NEB M0493L). PCR reactions were purified by PCR cleanup beads (UC Berkeley DNA Sequencing Facility) and eluted in 20 μL water. Half of the step-1 PCR was used as input for step-2 PCR with primers (UCSF CAT) containing unique dual indexes. Step-2 PCR reactions were performed in 50 μL as described above except with an annealing temperature of 72°C and 8-12 cycles, depending on the sample. The final libraries were purified with PCR cleanup beads and analyzed by Agilent Tapestation 2200. The libraries were pooled and sequenced on an Illumina Novaseq X instrument with single-end 100 bp reads (UCSF CAT).

### CiBER-seq screen processing and data analysis

Illumina sequencing reads were trimmed with cutadapt to obtain barcodes and UMIs^39^. Counts were extracted using the custom “bc-umi” program, which collapses highly similar barcodes and performs UMI de-duplication. These barcode counts were then collected into a matrix using the custom “bc-tabulate” program. Barcodes with an average count < 32 across all samples were discarded from downstream analysis. Barcodes were matched with sgRNAs in R using a custom script. Reads were analyzed using DESeq2, summing counts for each sgRNA across the independent barcodes. sgRNAs with an adjusted p-value < 0.01 were scored as statistically significant, and sgRNAs that also had a log_2_ fold change > 1.5 or < -1.5 were used for downstream GO analysis on PantherDB^31^.

### Individual validation of sgRNA-mediated phenotypes

Individual sgRNA expression vectors were cloned by first digesting a pCRISPRi-v3 expression base vector containing a GFP cassette with BstXI and BlpI and generating amplicons by PCR with overhangs containing each protospacer sequence and homology arms to the base vector. The protospacer sequences and primers used are listed in Table S2 and S4. Each sgRNA expression vector was packaged into lentiviruses in HEK293T Lenti-X cells and was individually transduced into cJK068 at an MOI < 1, resulting in ∼10% infected cells by GFP expression. Cells were sorted by GFP expression 4 days post transduction to select for sgRNA expression and recovered in media for 8 days. Cells were then induced with 5 ng/mL TNF-alpha for 2 hours in biological triplicate. Uninduced and induced cells were collected in TRI reagent for analysis of reporter expression and target gene knockdown efficiency by RT-qPCR using primers listed in Table S5.

## Data and code availability

Raw Illumina and PacBio sequencing data and processed barcode counts have been deposited in the NCBI Gene Expression Omnibus (GSE276055). Raw data and analysis scripts are available on Zenodo (https://doi.org/10.5281/zenodo.13687775).

---

**Supplementary Figure 1:**
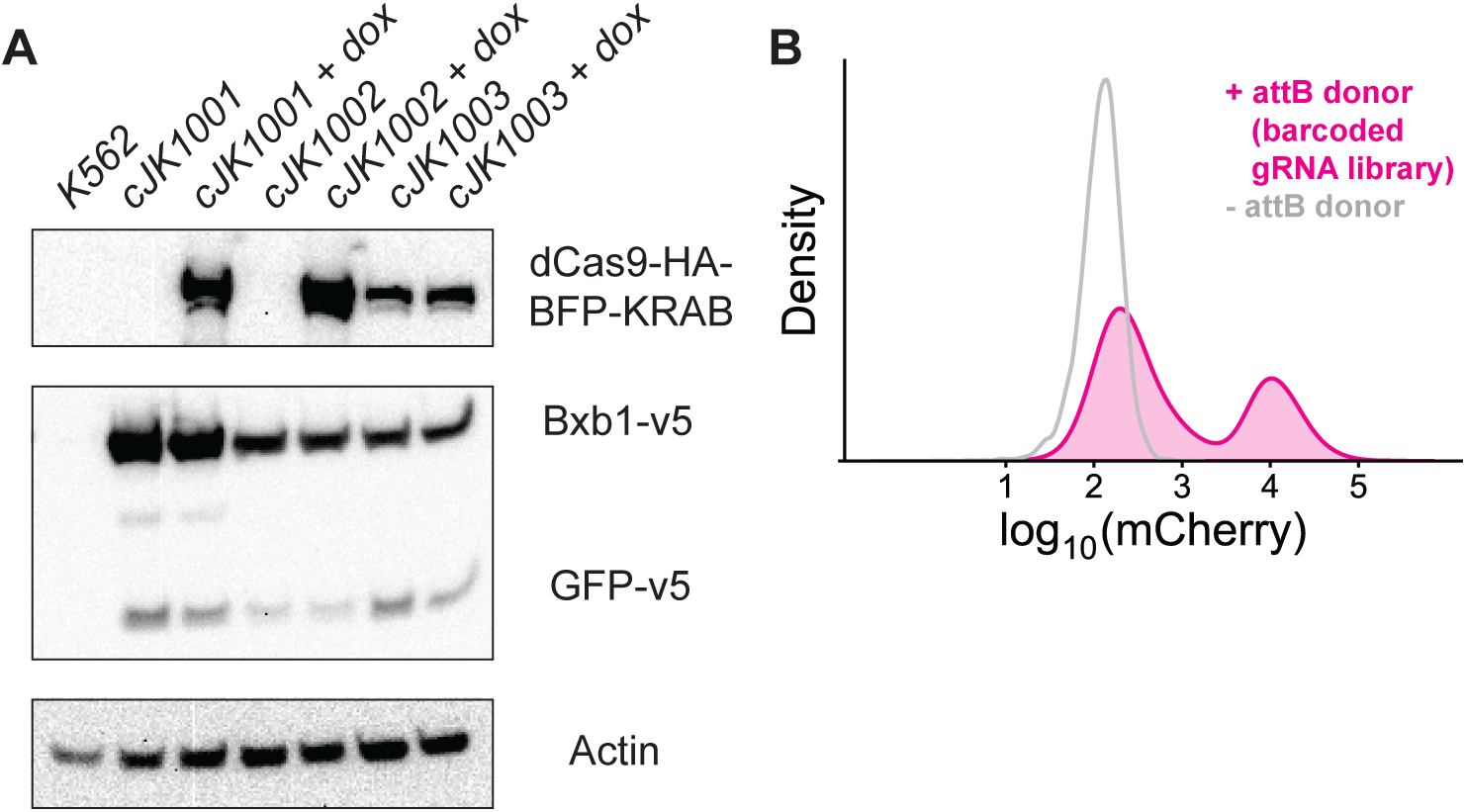
Further validation of mammalian CiBER-seq components. **(A)** Western Blot showing either inducible or constitutive dCas9-KRAB expression, Bxb1 or GFP expression, and actin as loading control. Cells were either left untreated or treated with 1 μg/mL doxycycline for 72 hrs. cJK1001: K562 cells with inducible dCas9-KRAB, two copies of attP landing pad. cJK1002: K562 cells with inducible dCas9-KRAB, one copy of attP landing pad. cJK1003: K562 cells with constitutive dCas9-KRAB, one copy of attP landing pad. **(B)** Cells that integrate the barcoded sgRNA library also express mCherry.

**Supplementary Figure 2:**
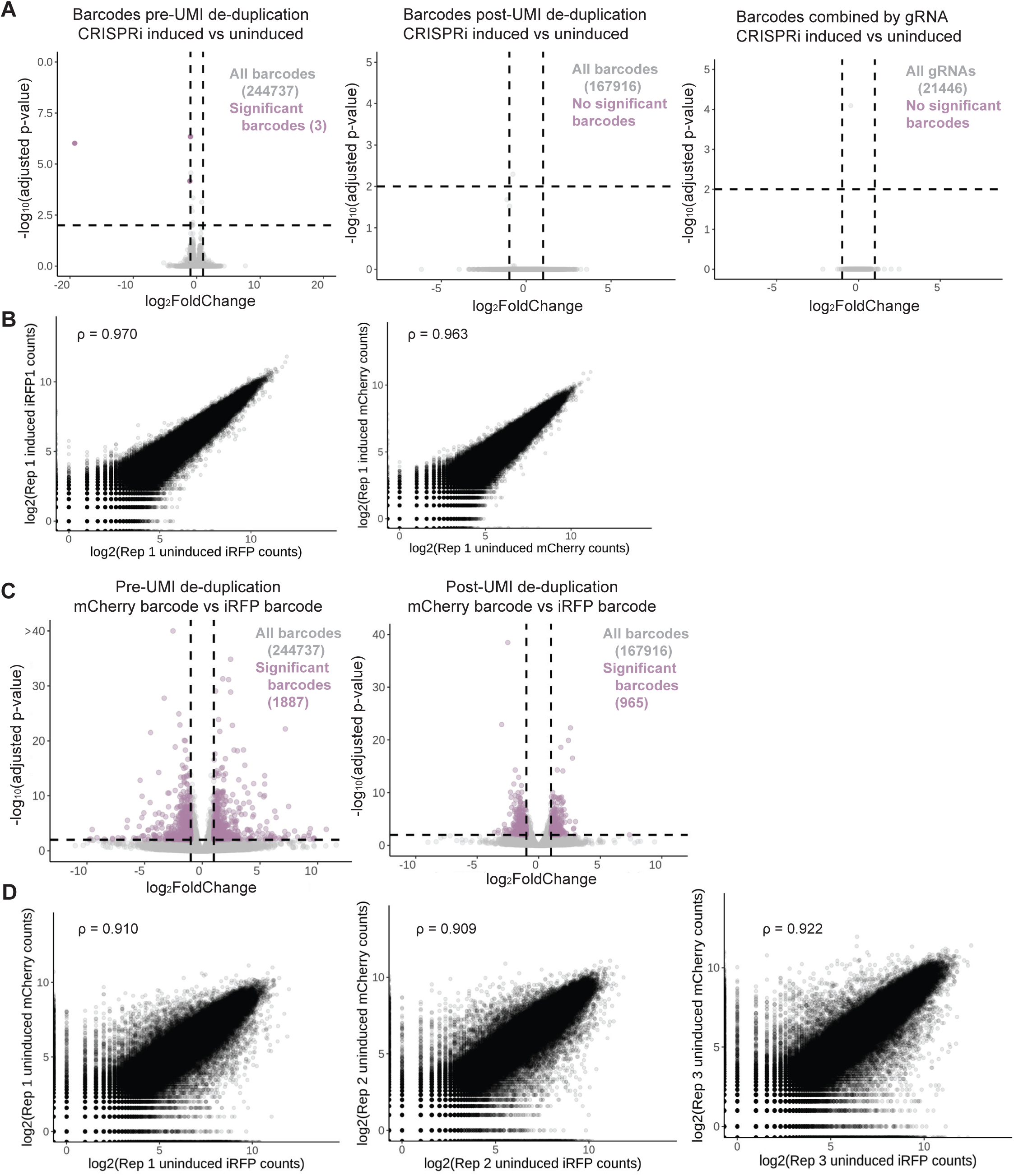
Additional validation of CiBER-seq screen for matched barcoded reporters. **(A)** DESeq2 comparison of CRISPRi induced vs uninduced samples for barcodes pre-UMI de-duplication, post-UMI de-duplication, and post combining barcodes by sgRNA. For the first two panels, each point represents one barcode, and for the last panel, each point represents one sgRNA. Dashed lines represent adjusted p-value < 0.01 and log2-fold change > 1 or < -1. **(B)** Comparison of either iRFP or mCherry barcode counts between uninduced and induced CRISPRi samples for replicate 1. Each point represents one barcode. **(C)** DESeq2 comparison of mCherry vs iRFP changes for barcodes pre-UMI de-duplication and post-UMI de-duplication (from both the pre- and post-CRISPRi induction samples). Each point represents one barcode. Dashed lines represent adjusted p-value < 0.01 and log2-fold change > 1 or < -1. **(D)** Comparison of iRFP and mCherry barcode counts for each replicate. Each point represents one barcode.

**Supplementary Figure 3:**
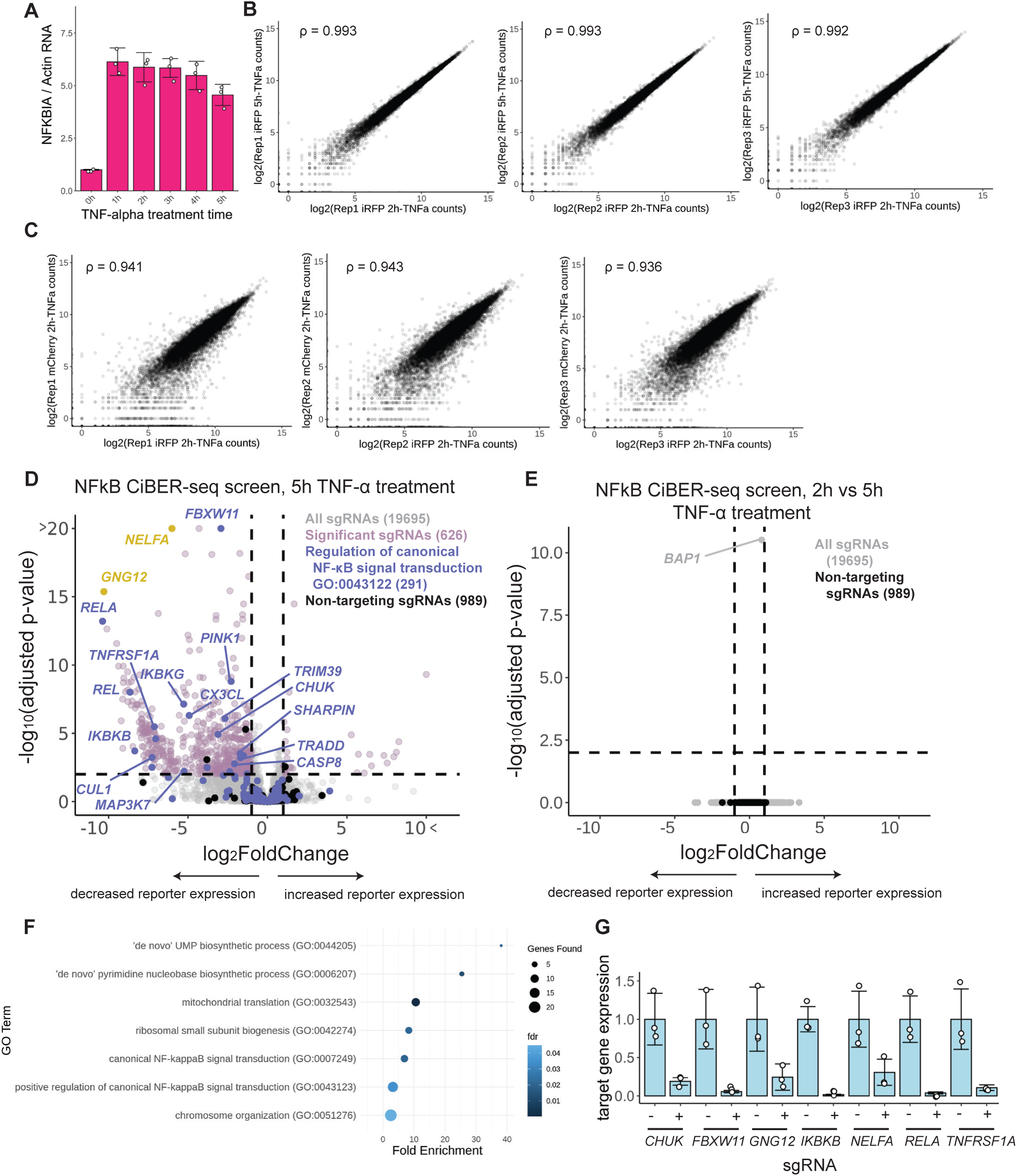
Additional validation of NF-κB CiBER-seq screen. **(A)** RT-qPCR of *NFKBIA* normalized to *ACTB* across different timepoints of TNF-alpha stimulation (n=3). **(B)** Comparison of iRFP barcode counts between the 2-hr and 5-hr timepoints. Each point represents an sgRNA. **(C)** Comparison of iRFP and mCherry barcode counts. Each point represents an sgRNA. **(D)** DESeq2 analysis as in Fig 3E. **(E)** DESeq2 analysis of genome-wide CiBER-seq screen for regulators of NF-κB, comparing the 2-hr and 5-hr timepoints. Each point represents a single sgRNA. Dashed lines represent adjusted p-value < 0.01 and log2-fold change > 1 or < -1. **(F)** Gene ontology statistical overrepresentation analysis of guides that prevent mCherry induction after 2 hrs of TNF-α treatment (log2FoldChange < -1 and adjusted p-value < 0.01). **(G)** RT-qPCR of each indicated endogenous gene normalized to *ACTB* in cells with and without the indicated, corresponding sgRNA (n=3).

## Notes

https://doi.org/10.5281/zenodo.13687775

https://www.ncbi.nlm.nih.gov/geo/query/acc.cgi?acc=GSE276055

